# Tibial nerve stimulation increases vaginal blood perfusion and bone mineral density and yield load in ovariectomized rat menopause model

**DOI:** 10.1101/2021.12.03.469332

**Authors:** Jiajie Jessica Xu, Lauren L. Zimmerman, Vanessa Soriano, Georgios Mentzelopoulos, Eric Kennedy, Elizabeth C. Bottorff, Chris Stephan, Kenneth Kozloff, Maureen J. Devlin, Tim M. Bruns

## Abstract

**Introduction and Hypothesis:** Human menopause transition and post-menopausal syndrome, driven by reduced ovarian activity and estrogen levels, are associated with an increased risk for symptoms including but not limited to sexual dysfunction, metabolic disease, and osteoporosis. Current treatments are limited in efficacy and may have adverse consequences, so investigation for additional treatment options is necessary. Previous studies have demonstrated that tibial nerve stimulation (TNS) or electro-acupuncture near the tibial nerve are minimally invasive treatments that increase vaginal blood perfusion or serum estrogen in the rat model. We hypothesized that TNS would protect against harmful reproductive and systemic changes associated with menopause.

**Methods:** We examined the effects of twice weekly TNS (0.2 ms pulse width, 20 Hz, 2x motor threshold) under ketamine-xylazine anesthesia in ovariectomized (OVX) female Sprague Dawley rats on menopause-associated physiological parameters including serum estradiol, body weight, blood glucose, bone health, and vaginal blood flow. Rats were split into three groups (n = 10 per group): 1) intact control (no stimulation), 2) OVX control (no stimulation), and 3) OVX stimulation (treatment group).

**Results:** TNS did not affect serum estradiol levels, body weight, or blood glucose. TNS transiently increased vaginal blood perfusion during stimulation for up to 5 weeks after OVX and increased areal bone mineral density and yield load of the right femur (side of stimulation) compared to the unstimulated OVX control.

**Conclusion:** TNS may ameliorate some symptoms associated with menopause. Additional studies to elucidate the full potential of TNS on menopause-associated symptoms under different experimental conditions are warranted.

**Summary:** Percutaneous tibial nerve stimulation increases vaginal blood perfusion, areal bone mineral density, and femur yield load in an ovariectomized rat model of menopause.

## INTRODUCTION

Menopause, defined as the cessation of menstrual periods for at least 12 consecutive months [1] and characterized by low levels of ovarian hormones (17β-estradiol and progesterone) [2] in humans, is part of reproductive aging. Approximately 6,000 women in the United States reach menopause every day [1], and 4 out of 5 women will experience psychological or physical symptoms around menopause [3].

Traditional vasomotor symptoms and related psychosocial impairment during the menopausal transition (hot flushes, night sweats, sleep disturbances, mood swings, cognitive deficits, social impairment, and more) can lead to a decreased quality of life in perimenopausal women [1, 4]. In addition the menopause transition is associated with an increase in health risk factors including but not limited to sexual dysfunction, cardiovascular disease, osteoporosis, cancer, diabetes, and stroke [2]. While hormone therapy remains the most effective treatment for vasomotor motor and genitourinary symptoms of menopause and has been shown to prevent bone loss and fracture, risks of hormone therapy include coronary heart disease, stroke, venous thromboembolism, and dementia [5].

An alternative treatment method may be tibial nerve stimulation. Tibial nerve stimulation has been used clinically to treat pelvic organ disorders such as overactive bladder, fecal incontinence, and pelvic pain [6]. The most common application of tibial nerve stimulation, for the treatment of overactive bladder, has equivalent or superior effects and fewer side effects (dry mouth, constipation) compared to an existing pharmacologic treatment (anti-cholinergic medication). The clinical procedure is minimally invasive and has minimal side effects [6].

While the use of tibial nerve stimulation for pelvic disorders is promising, the exact mechanism for how the treatment works is unclear [6, 7]. Tibial nerve stimulation is a form of neuromodulation (specifically electrical neural stimulation), which works by stimulating or blocking the flow of action potentials through the nervous system [8]. In tibial nerve stimulation, it is believed that stimulation of somatic afferent nerves that project to the lumbosacral spinal cord leads to the release of excitatory neurotransmitters that activate ascending pathways to the brain or spinal circuits, which modulate visceral sensory and involuntary motor mechanism [7].

Previous work in our laboratory shows that tibial nerve stimulation has potential for treatment of sexual dysfunction symptoms. We demonstrated that tibial nerve stimulation increases vaginal blood perfusion (an indicator of sexual arousal in humans [11]), in intact anesthetized rats [9], and can also improve sexual dysfunction symptoms in humans (pilot study) [10]. However, these studies did not include a control group. In addition, preliminary as-yet unpublished data from our laboratory shows tibial nerve stimulation increases sexual receptivity in ovariectomized (OVX) rats [12]. Similar studies using electroacupuncture at the “Sanyinjiao (SP6)” acupoint (near the ankle, around the site where the tibial nerve is percutaneously accessed) have improved perimenopausal symptoms and increased serum or ovarian estradiol levels in rats [13] and humans [14]. Additional electro-acupuncture studies at other points in conjunction with the SP6 acupoint have increased serum estradiol levels specifically in OVX rats [15, 16].

In general, these studies are limited to the direct effects of tibial nerve stimulation or SP6 electroacupuncture on the reproductive system (estradiol levels, vaginal blood perfusion, sexual dysfunction or perimenopausal symptoms). To our knowledge, no study has examined the effects of tibial nerve stimulation or SP6 electro-acupuncture on the systemic effects of menopause. For instance, menopause-associated estrogen decline predisposes women to osteoporosis (resulting decreased bone mineral density and changes in bone biomechanical properties such as yield load), which leads to an increased risk for osteoporotic fractures and is considered one of the largest public health priorities by the World Health Organization [17]. In this study, we examined the effects of chronic (twice a week for 6 weeks) percutaneous tibial nerve stimulation (PTNS) in OVX rats. We hypothesized that in addition to improving reproductive parameters (serum estradiol, vaginal blood perfusion), the increase in serum estradiol would protect against some harmful systemic changes of menopause (ex: weight gain, osteoporosis, diabetes).

## METHODS

All experimental procedures were performed at an AAALAC accredited institution (Protocol #: PRO00009736), according to the standards established by the Guide for the Care and Use of Laboratory Animals and were approved by the University of Michigan Institutional Animal Care and Use Committee.

### Animals

Female CD^®^ (Sprague Dawley) IGS rats (250-300 g [11-15 weeks], n = 30, Charles River Breeding Labs, Wilmington, MA) were used in this study and acclimated for a week prior to surgical manipulation (ovariectomy [OVX] or sham procedure). Sprague Dawley rats are a standard preclinical model for menopausal research, and have been used to examine the effects of ovariectomy on cardiovascular function, cancer, bone, and cognition [18]. Rats were split into 3 groups (n=10 per group, consistent with previous similar studies examining the effects of stimulation on reproductive parameters [15]): 1) intact control, 2) OVX control, and 3) OVX stim.

The OVX control and OVX stim groups underwent an OVX procedure, while the intact control group underwent a sham procedure. Only the OVX stim group received PTNS. After surgery, the rats were rested for 2 weeks, during which vaginal cytology was used to establish the presence of normal estrous cycling in intact rats and cessation of cycling in OVX rats. The OVX stim rats were stimulated twice a week for 6 weeks, and blood and vaginal cytology were collected once a week during this time until a terminal necropsy (Figure 1). Clinical PTNS in humans is performed once a week up to 12 weeks [19], so in this study we sought to do 12 sessions, but at an increased frequency because rats have a shorter estrus cycle.

**Figure 1:**
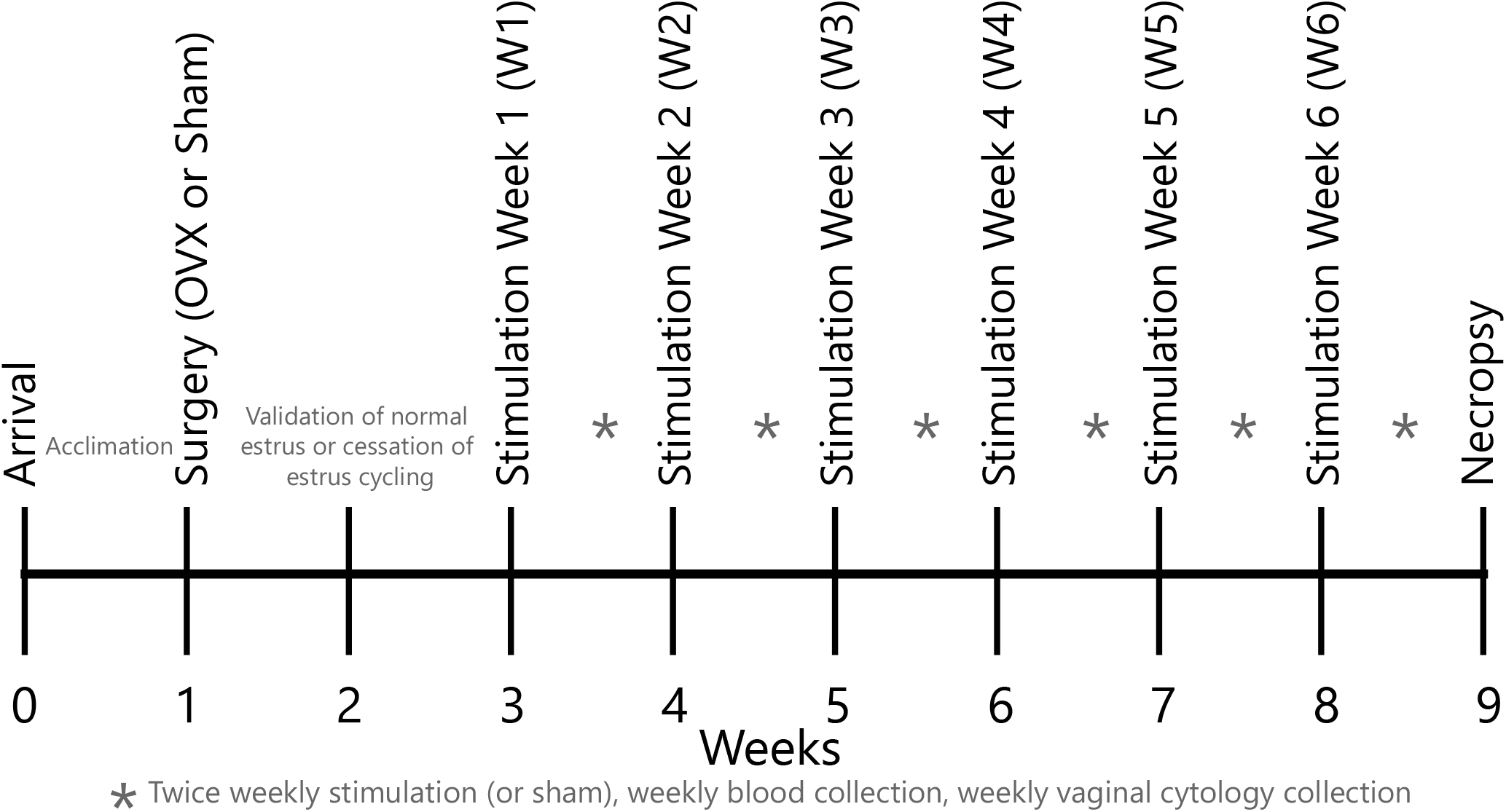
Experimental timeline of study. OVX = ovariectomy.

### Ovariectomy (OVX) or sham surgery

Anesthesia and analgesia for OVX and sham procedures: For all groups, animals were anesthetized with isoflurane anesthesia (1-5%). Reflexes were checked for appropriate depth of anesthesia prior to surgical incision. Carprofen (5 mg/kg, Rimadyl, Zoetis Animal Health, Parsippany-Troy Hills, NJ) was given subcutaneously immediately before surgery, as well as the following day. The animals were monitored daily until wound clips were removed, 7 days after surgery.

OVX (n=20) and sham (n=10) procedures: The OVX and sham procedures were performed consistent with previously described methods (Appendix S1).

### Anesthesia for test (stimulation or control/sham) sessions

Ketamine-xylazine anesthesia (50-70 mg/kg ketamine [Zetamine, Vet One, Boise, ID], 5-7 mg/kg xylazine [AnaSed, Akorn, Lake Forest, IL]) was induced by intraperitoneal injection, twice weekly, for 6 weeks in all groups of rats. Ketamine-xylazine has been used in previous studies evaluating sexual arousal in sedated rats [20, 21], and provides greater sedation than ketamine alone.

Rats that did not reach adequate anesthetic depth on their given day were allowed to recover and were re-dosed the following day. Rats were kept on a warming pad during the duration of anesthesia. The anesthesia was reversed with atipamezole (0.25 mg per rat, Antisedan, Orion Corporation, Espoo, Finland) subcutaneously. Regardless of experimental group, rats were anesthetized for 45-60 minutes before reversal. The anesthetic regime was modified during the study due to anesthetic-associated animal death and delayed recovery early in the study (Appendix S2).

### Percutaneous Tibial Nerve Stimulation

For the OVX stim group only, during the twice weekly anesthesia sessions, an EMG wire pair (EMT-2-30 subcutaneous bipolar hook electrode, MicroProbes for Life Science, Gaithersburg, MD) was inserted subcutaneously near the ankle, and connected to an isolated pulse generator (Model 2100 Isolated Pulse Stimulator, A-M Systems, Sequim, WA). The amplitude of stimulation (biphasic, rectangular, 0.2 ms pulse width, 20 Hz) was increased until the hind paw motor threshold was detected. The tibial nerve was then stimulated at twice the motor threshold continuously for 30 minutes, preferentially on the right side (Appendix S3). OVX control and intact control groups did not receive EMG wire placement or stimulation during anesthesia. The EMG wire was not placed, and stimulation was not performed for the OVX control or intact control groups.

### Antemortem measures

#### Weight

Rats were weighed at baseline (before surgery), once a week during the 6-week stimulation period, and once at necropsy. The normalized weight was calculated for each rat at each time point by dividing its current weight by its baseline weight.

#### Vaginal blood perfusion (VBP)

VBP during anesthesia sessions was assessed using a laser Doppler probe (MNP110XP, ADInstruments, Colorado Springs, CO, USA) connected to a Blood FlowMeter (50 Hz sampling rate; INL191, ADInstruments) as previously described [9, 22]. The base of the probe was secured to the tail using tape to prevent the probe from sliding out, as rats often urinated during anesthesia. VBP was sampled for 5 minutes before stimulation (5-minute baseline) and 30 minutes during stimulation (sham stimulation in the OVX control and intact control groups) in each rat during weeks 1, 3, and 6.

VBP signals were analyzed using a time frequency representation with a continuous transform method in MATLAB (Mathworks, Natick, MA, USA). The neurogenic band (0.076-0.200 Hz) was isolated, and scalogram energies calculated using a previously described approach [9, 22]. The scalogram energy during the stimulation period was divided by the average of the 5-minute baseline period to generate a continuous normalized neurogenic energy trace for the entire stimulation period, as well as three 10-minute normalized values per recording session (0-10 min, 10-20 min, 20-30 min). A positive response was defined as at least a 500% increase neurogenic energy compared to baseline [9]. The normalized neurogenic energy values for each group (separated by week and 10-minute timepoint interval) were compared to a hypothetical normalized mean of 1 (indicating no change) using a post-hoc one-sided Wilcoxon signed-rank test.

#### Blood collection for blood glucose and serum estradiol

Once weekly, unanesthetized blood glucose was measured with a glucometer (AlphaTrak2, Zoetis, Parsippany-Troy Hills, NJ) by pricking the tail tip or a tail vein. Under anesthesia, 1 ml of blood (< 0.5% body weight) was collected from each rat using the sublingual vein during recovery from the stimulation or sham stimulation. Serum was extracted from whole blood, and serum estradiol was analyzed using a commercial radioimmunoassay kit with a double-antibody method (Catalog # DSL-4800, Diagnostic Systems Labs [DSL], Webster, TX, USA).

#### Vaginal cytology

Once weekly, coinciding with anesthesia sessions, vaginal cytology was obtained by flushing 0.5 to 1 ml of sterile saline into the vaginal canal with a bulb pipette, and examining the sample under a microscope. The estrous phase was determined by assessing cell morphology [23]. Select samples (weeks 3, 4, 6) were further quantified (% leukocytes, cornified cells, and nucleated epithelial cells), with Adobe Photoshop (Adobe, San Jose, CA, USA) using counting, brighten, and sharpen features.

### Postmortem measures

Postmortem samples from one animal (OVX control) that died during a sedated session in the second half of week 6 was included in the necropsy data. Postmortem samples from other animals that died previously were not included (specific animals detailed at start of Results).

#### Gross necropsy

Animals were euthanized with carbon dioxide 3 days after the end of the 6^th^ week and necropsied immediately afterwards. The final rat weights were obtained through a different scale than previously used for during antemortem weights, and 3-5 ml blood was collected through a cardiac puncture for serum estradiol analysis. The heart, uterus, tibial nerves (right and left), bladder, and femurs were collected. The wet weight of the uterus was measured at the time of necropsy after being separated from attached organs, and trimmed at the oviduct and immediately below the cervix as previously described [24]. Cross sectional areas of both the right and left uterine horn were taken at the point of uterine bifurcation. Due to grossly unremarkable findings of other organs, only the uterus and tibial nerves were evaluated histologically.

#### Histological analysis

The nerves and uterus were stained using H&E. The nerves became separated and distorted during processing, so no further evaluation was performed (although no baseline pathology was noted on brief examination). The uterine cross-sectional area (excluding inner luminal area) was measured using QuPath (https://qupath.github.io/) as a measure of estrogenic effect using the magic wand tool.

#### Femur analysis

The femurs were submitted for dual-energy x-ray absorptiometry (DXA) scanning, micro-computed tomography (micro-CT), and biomechanical testing. In the time between necropsy and additional testing, the femurs were wrapped in saline-soaked gauze and stored at −80 °C. For DXA scanning, the right and left femurs were scanned *ex vivo* on an excised bone positioner plate (pDXA, PIXImus I, GE Lunar Corp., Madison, WI, USA) to measure areal bone mineral density (aBMD) [25]. For micro-CT, the right femurs were scanned in water using cone beam micro-computed tomography (Skyscan 1176, Bruker, Billerica, MA) (Appendix S4). Output parameters included trabecular properties (bone mineral density, bone volume fraction, trabecular thickness, trabecular number) and cortical properties (tissue mineral density, total area, bone area, cortical thickness, polar moment of inertia, and bone marrow area). Biomechanical properties of the right femur (three-point bending, mid-femoral diaphysis) were tested on a material testing machine (MTS Systems, Eden Prairie, MN, USA). Stiffness, yield load, ultimate load, fail load, yield displacement, ultimate displacement, fail displacement, total work, and post yield displacement were calculated as previously described [26].

### Data Analysis

An Analysis of Variance (ANOVA) test was used to determine if differences existed between groups. For most measures, pairwise comparisons between groups were calculated using JMP Pro 16 (SAS Institute Inc, Cary, NC, USA) using the Tukey-Kramer HSD test (parametric assumption, adjusting for multiple comparisons). For VBF and vaginal cytology, the data was non-parametric, so it was analyzed using the Steel Dwaas test (non-parametric equivalent of Tukey-Kramer HSD). Differences were significant for p < 0.05. Where relevant, data is reported as mean ± standard deviation. Due to the nature of the study, experimenters could not be blinded to the experimental groups, however, analysis of DXA, micro-CT, and biomechanical testing was done in collaboration with individuals who were blinded to the different treatment groups.

## RESULTS

Of the original thirty animals (ten animals per group, three groups), seven animals (23%) were lost during the study (Appendix S5). At necropsy, twenty-three animals remained (intact control [n = 9], OVX control [n = 7], OVX stim [n = 7]).

In context of twice a week anesthesia sessions, we observed 6 deaths out of more than 300 anesthetic events (2% rate), and the survival rate improved significantly after the addition of reversal agents and cessation of re-dosing. The changes made in anesthetic protocols to improve animal welfare were made across all groups simultaneously, and thus unlikely to result in a bias towards any experimental group. Despite this, it is noted that the OVX groups suffered a greater mortality rate than the intact control group. This may be due to differences in physiology after the OVX surgery.

### Antemortem measures

#### Weight

All groups generally gained weight or maintained weight over the course of the study when tracked within group over time (except for slight decreases in weight in week 2 and 4 of the OVX control group) (Figure 2). During necropsy, all groups apparently lost weight compared to the previous week, but this may be due to the use of a different scale during the necropsy measurement, and not a true decrease in weight. In general, weights of both the OVX control and the OVX stim groups were significantly increased at each time point compared to intact control, with few exceptions. There was no significant difference between OVX control and OVX stim at any timepoint (Table S1).

**Figure 2:**
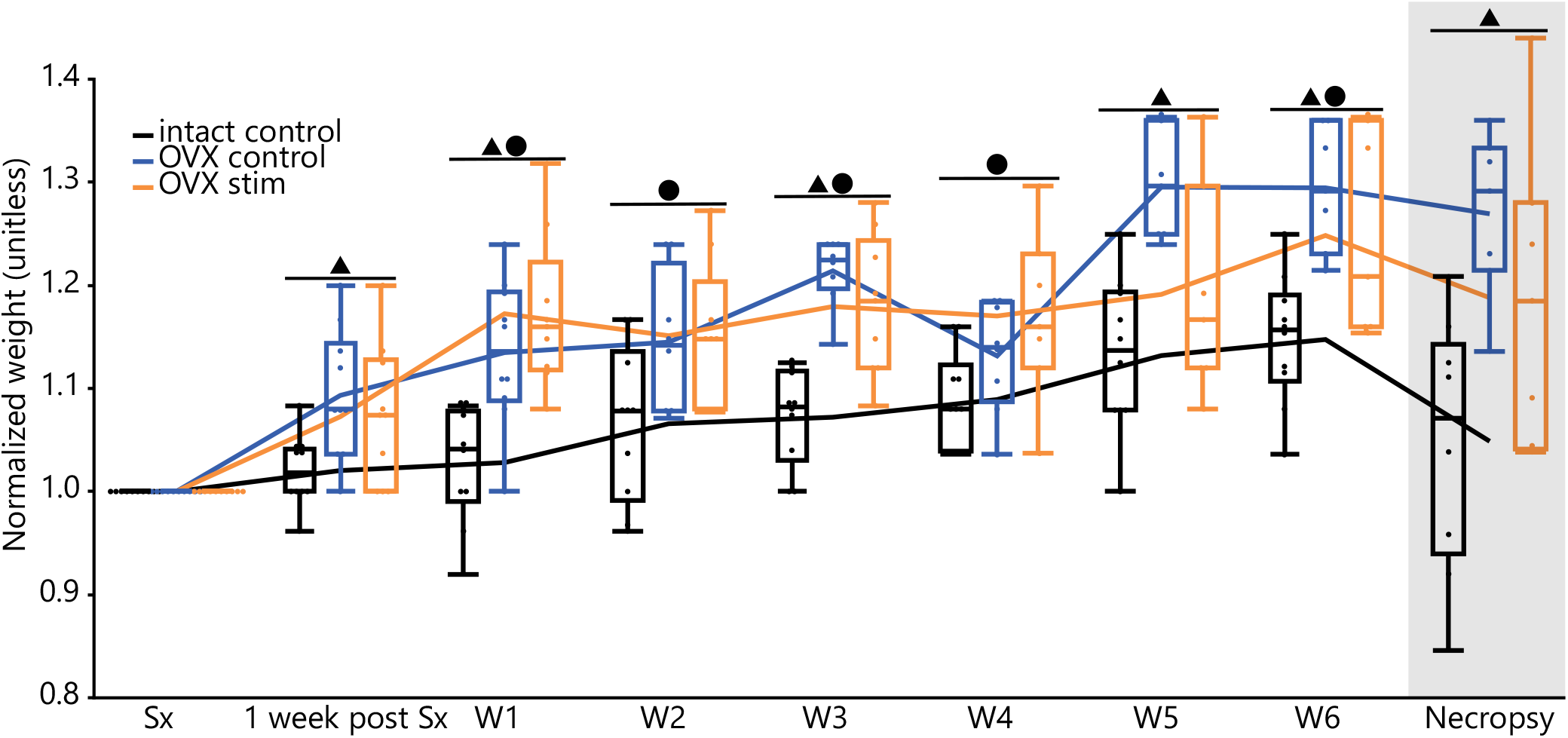
Box plots of normalized weight over time grouped by timepoint (black/left = intact control, blue/middle = OVX control, orange/right = OVX stim for each timepoint). Small dots represent individual measures. Boxes represent 1st to 3rd quartile range (interquartile range [IQR]), with the line in the middle of each box representing the median. Tips of whiskers extending below and above boxes represent 1st quartile - 1.5*IQR and 3rd quartile + 1.5*IQR respectively. Lines connect means across time points. Shaded area represents samples taken at necropsy with a different weigh scale configuration. ▲ denotes significant difference (p < 0.05) between OVX control and intact control. ● denotes significant difference (p < 0.05) between OVX stim and intact control.

#### VBF

Overall, PTNS appeared to increase VBF compared to baseline. The OVX stim group had both a greater duration and amplitude of increase in VBF after stimulation compared to the other unstimulated groups, as well as a higher rate of response to stimulation (positive response defined as > 500% increase in neurogenic energy compared to baseline) (Figure 3). In the OVX stim group, 67% of animals responded during week 1 (at both 10-20 min and 20-30 min), and 63% of animals responded during at week 3 (20-30 min). In all other groups and timepoints, the response rate was < 50% (0-44%).

**Figure 3:**
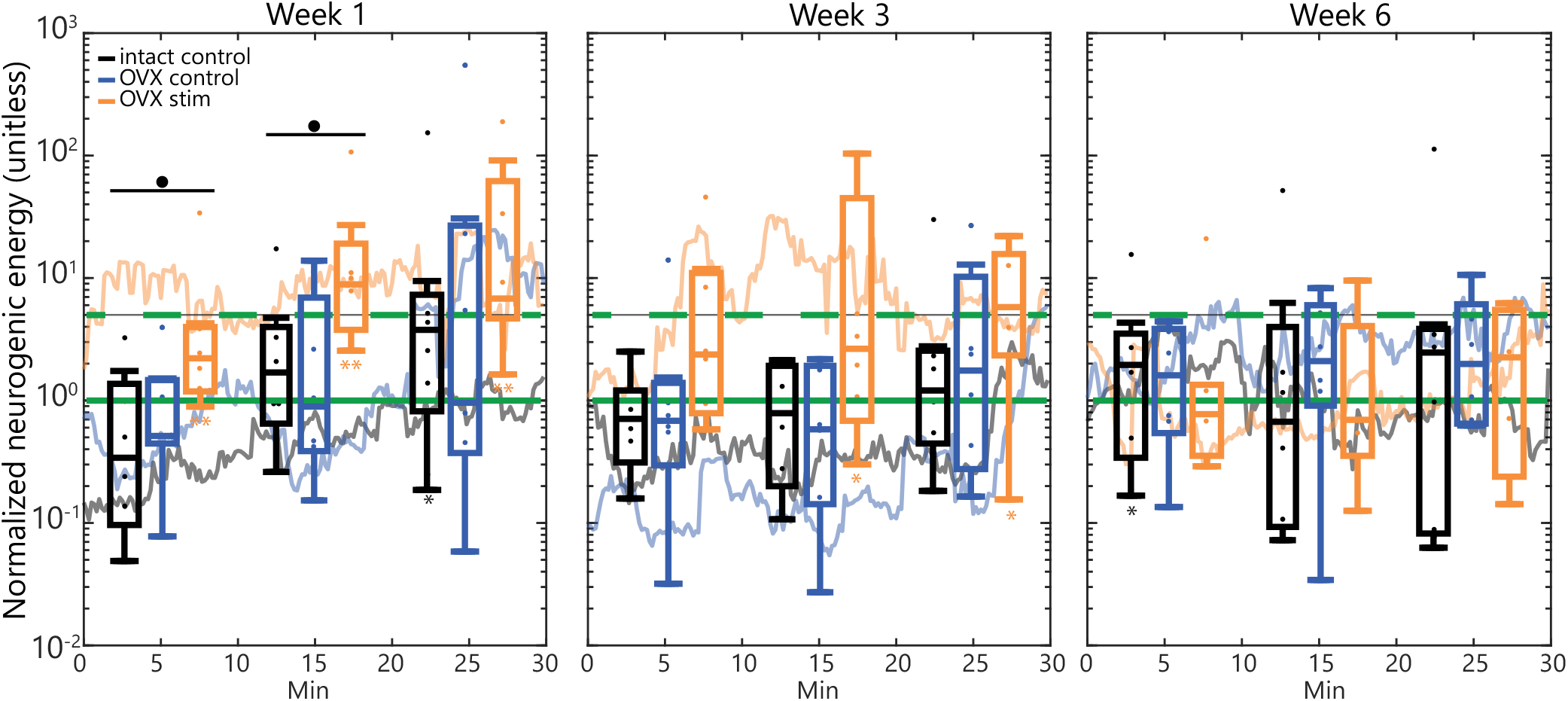
Mean value trace (signal averaged every 10-seconds) and box plots (for pooled 10-minute average values) representing the vaginal blood perfusion (expressed as normalized neurogenic energy) of all rats during 30-minute stimulation or sham stimulation periods. Box plots are organized as in Figure 2. Dashed green line denotes 5x baseline value, and solid green line denotes 1x baseline value. ● denotes significant difference (p < 0.05) between OVX stim and intact control. Asterisks beneath each box plot denote significant increase (* p < 0.05, ** p < 0.01) from baseline of session (normalized neurogenic value = 1).

Using a post-hoc one-sided Wilcoxon signed-rank test, the OVX stim group was significantly increased from baseline at the most timepoints and at greater levels of significance compared to the unstimulated groups (intact control, OVX control). The intact control group was also significantly increased from baseline at several timepoints. The OVX control group was not significantly increased from baseline at any time (Table S2).

The OVX stim group had significantly increased normalized neurogenic energy in the VBF measures compared to the intact control group during the 0-20 minutes of stimulation during week 1 (Figure 3, Table S3). The peak VBF increase in the OVX stim group was seen at 20-30 minutes during week 1 and at 10-20 minutes during week 3 (Figure 3).

#### Blood glucose, serum estradiol, vaginal cytology

Blood glucose did not differ between groups. Serum estradiol and vaginal cytology differed as expected between intact and OVX animals (increased serum estradiol and percent cornified cells in intact group). No significant difference was observed between the OVX stim and OVX control groups, which would suggest an effect from stimulation (Appendix S6, Table S4).

### Postmortem measures

#### Gross necropsy

At necropsy, uterine wet weights were significantly higher (p < 0.0001) in the intact control group (0.8 ± 0.3 g) compared to the OVX control (0.1 ± 0.02 g) and OVX stim (0.2 ± 0.04 g) groups. The uterine wet weights were not significantly different between the two OVX groups.

#### Histological analysis

The cross-sectional area of the uterus was significantly higher (p < 0.0001) in the intact control group (8,087,360 ± 2,548,981 μm^2^) compared to the OVX control (1,535,158 ± 400,171 μm^2^) and OVX stim groups (1,786,237 ± 487,604 μm^2^). There was no significant difference between the two OVX groups.

#### Femur analysis

DXA analysis showed aBMD was significantly higher in the intact control group as compared to the OVX control on both the right and left side in DXA scanning (Figure 5, Table S5, S8). The OVX stim group had a significantly higher aBMD compared to OVX control on the right side (Table S5, S8). Micro-CT analysis of the right femur showed that compared to the intact control rats, both OVX groups had significantly lower trabecular BMD, bone volume fraction (BV/TV), trabecular number, and higher trabecular thickness (Table S6, S8). There were no differences between any groups in any other measured outcome (Table S8). Biomechanical analysis of the right femur showed yield load of the right femur was significantly higher in the OVX stim group as compared to the OVX control and was not different from the intact control in the biomechanical testing (Figure 6). There were no significant differences between any groups in other biomechanical parameters (ultimate load, stiffness, total work, yield displacement, ultimate displacement, fail load, fail displacement, or post yield displacement) (Tables S7, S8).

## DISCUSSION

In this study we examined the effects of a standard clinical treatment for pelvic organ dysfunction, PTNS, on serum estradiol and menopause-associated symptoms in a rat model of menopause. We hypothesized that PTNS would increase serum estradiol levels in OVX rats compared to unstimulated OVX controls, leading to changes in other post-menopausal parameters that would approach measures for intact control rats.

In contrast to previous human and animal electro-acupuncture studies where stimulation of the “SP6” acupoint has increased estradiol levels [13, 14], PTNS in this study did not affect serum estradiol levels in OVX rats (Figure 4). Therefore, unsurprisingly, many other menopause-associated parameters expected to be driven by serum estradiol also did not differ significantly between the OVX stim and OVX control groups.

**Figure 4:**
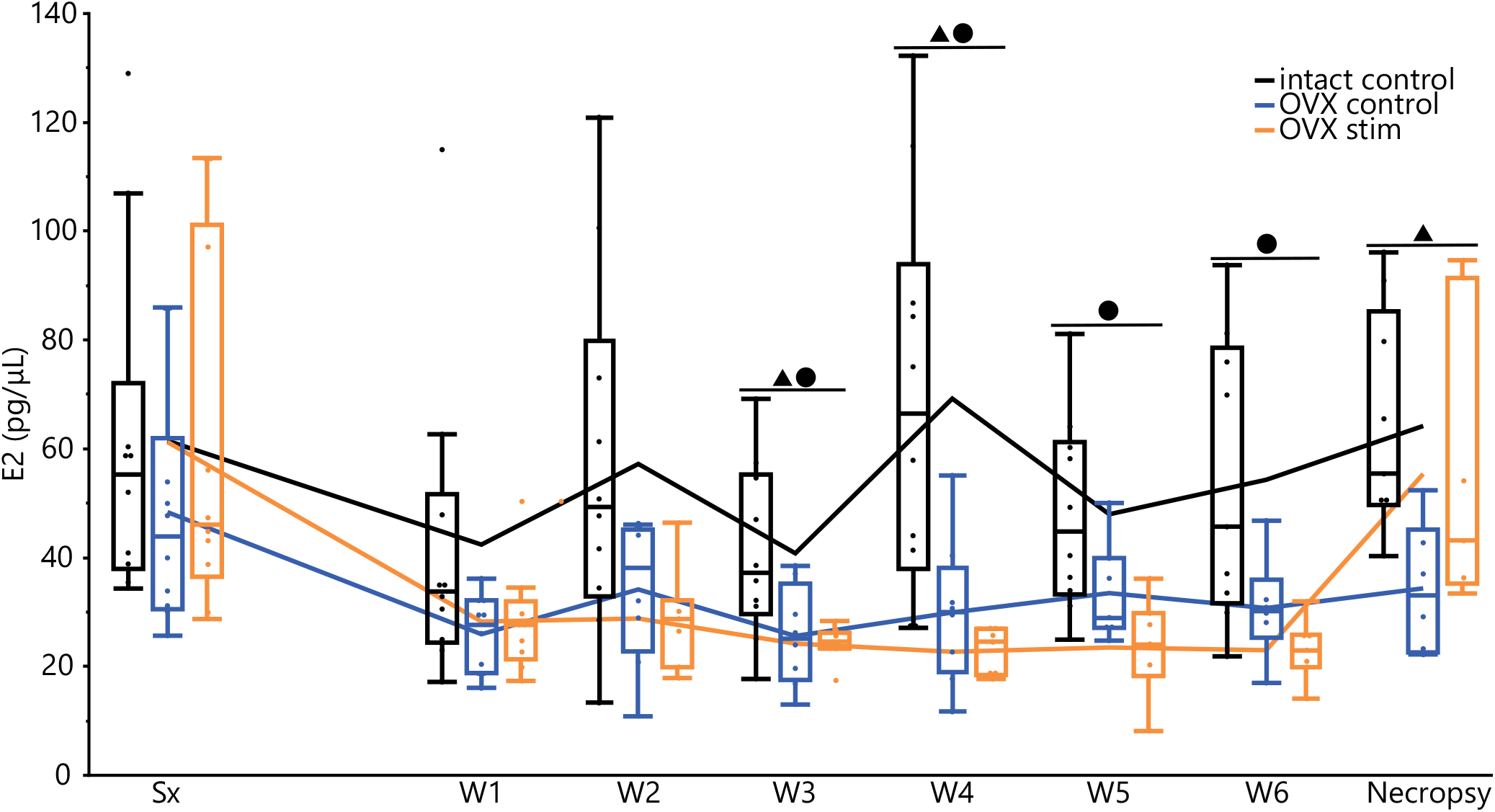
Box plots of estradiol (E2, pg/μL) per experiment, organized as in Figure 2. Lines connect means across time points. ▲ denotes significant difference (p < 0.05) between OVX control vs intact control, ● denotes significant difference (p < 0.05) between OVX stim and intact control).

However, we found that PTNS transiently increased vaginal blood perfusion (VBP) for up to 5 weeks after OVX surgery (Figure 3). Eight weeks after OVX, VBF ceased to be affected by PTNS, however. The OVX surgery causes vaginal atrophy [27] and decreases vaginal innervation [28] in both rats and humans [29]. Over time, this could decrease the target organ’s ability to be stimulated or provide an adequate response to PTNS. During week 1, VBP in the OVX stim group peaked around 20-30 minutes after the onset of stimulation, consistent with previous terminal studies in intact rats [9]. However, during week 3, VBP peaked earlier, at around 10-20 minutes. This may be due to either differences between the OVX and intact rat model or neural habituation to the repeated PTNS treatment. During peak response times in the OVX stim group, the response rate of animals that exceeded a threshold of 5x baseline VBF (63-67%) was similar to previous studies (75.8%) [9]. PTNS is a potential alternative to hormone replacement therapy in treating genitourinary symptoms such as vaginal dryness during the menopause transition period. However, it is unclear if this effect can be sustained over time.

PTNS appeared to partially rescue OVX-induced bone loss on the stimulated side. The OVX stim group had higher aBMD compared to the OVX control group in the right femur (generally the stimulated side) as measured by DXA (Figure 5, Table S8). The OVX stim group also had a higher yield load in the right femur compared to the OVX control group with biomechanical testing (Figure 6, Table S8). Interestingly, there were no differences between OVX control and OVX stim on the left (typically unstimulated) side. These results suggest that PTNS may have localized beneficial effects in menopausal patients who are at risk of osteoporosis.

**Figure 5:**
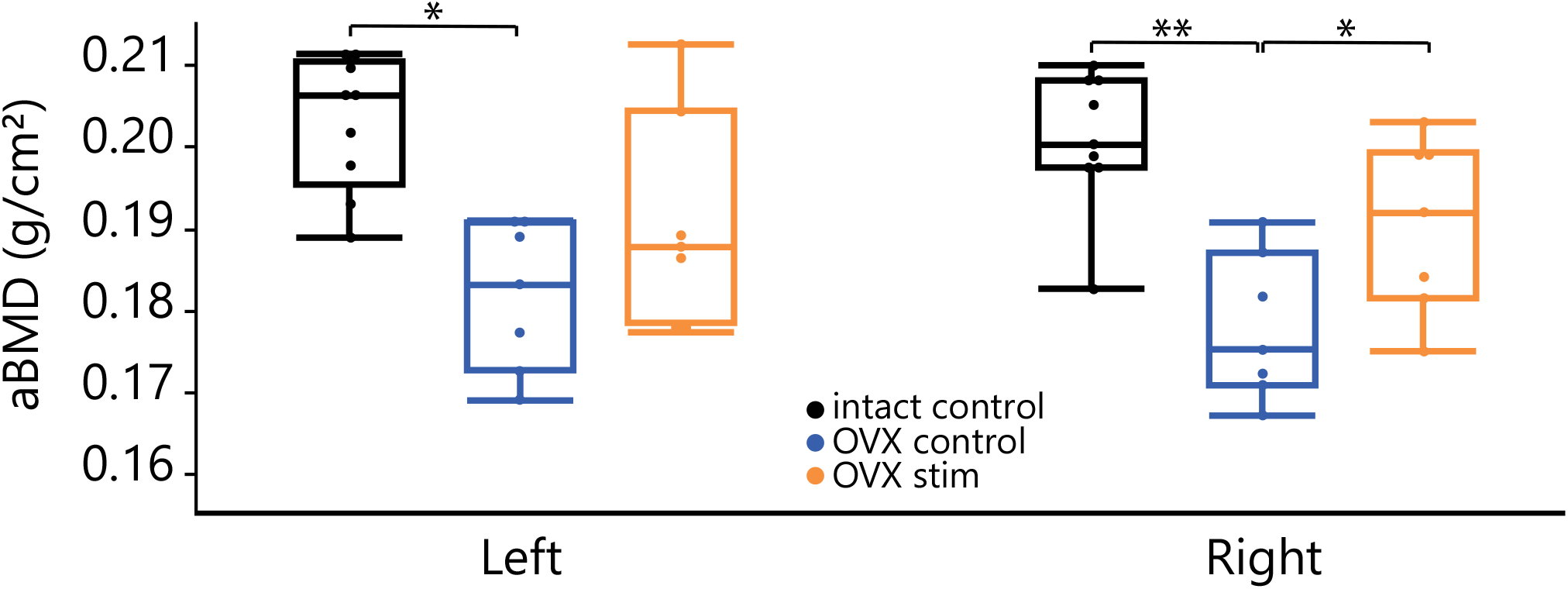
Box plots of areal bone mineral density (aBMD) for each group, separated by limb side (left vs right), as organized in Figure 2. Stimulation was primarily on the right side. Points represent individual samples. * p < 0.05, ** p < 0.01.

**Figure 6:**
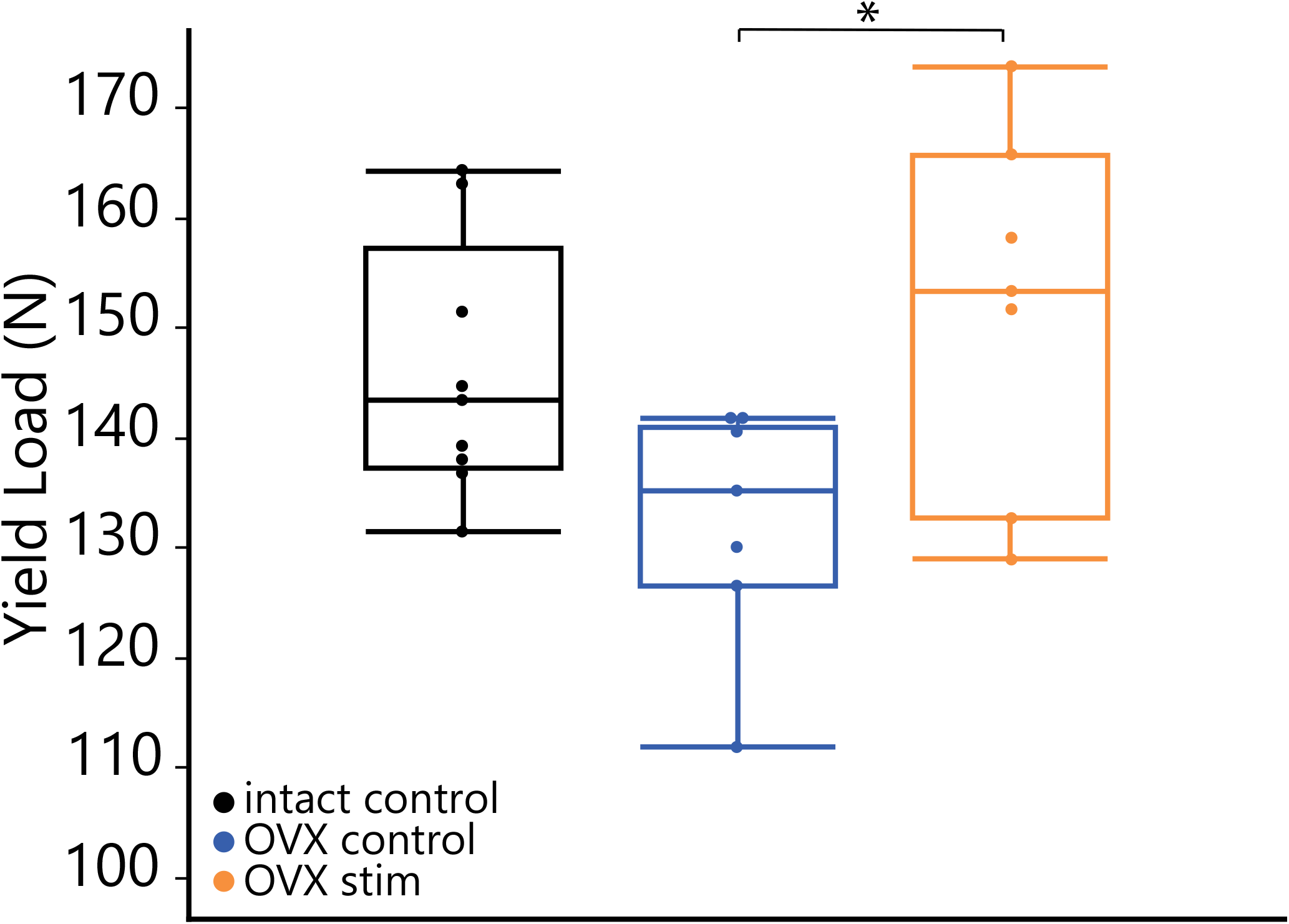
Box plots of yield load for each group, for right femur (the primary site of stimulation), as organized in Figure 2. Points represent individual samples. * p < 0.05.

Stimulation parameters in this study differed from a previous electro-acupuncture rat study in several ways [13]. 1) Stimulation frequency in this study was 20 Hz (compared to alternating between 2 and 100 Hz). 2) Stimulation frequency across sessions was twice a week in this study (compared to every other day). 3) Stimulation duration during per session in this study was 30 minutes (in contrast to 20-minute duration). 4) Stimulation was unilateral (compared to bilateral). The parameters selected in this study were consistent with previous nerve stimulation studies (tibial [9] or pudendal [22, 30] nerves) in rats and PTNS in humans [10]. It is possible that changing stimulation frequency could affect output, as demonstrated by the frequency-dependent effects of PTNS on the bladder (6-20 Hz is inhibitory, while 2 Hz is excitatory) [31]. Stimulation more frequently than twice a week may lead to greater or faster improvement, as suggested by clinical studies for fecal incontinence [32] and overactive bladder [33]. It is possible that bilateral PTNS could increase treatment efficacy; bilateral tibial nerve stimulation has been shown to be effective for constipation [34, 35] and overactive bladder [36].

Several other studies demonstrating the effects of electro-acupuncture on extragonadal aromatization of estradiol in an OVX rat model [15, 16] have also included additional acupoints along the dorsal and abdominal midline. While it has been demonstrated that the ankle PTNS access site or acupoint “SP6” alone should still significantly increase serum estradiol level [13, 14], the additional points may enhance estradiol production, and are worth investigating in future studies. The exact mechanisms of these additional points are unknown, but there are several proposed hypotheses (Appendix S7). Based on the many similarities between PTNS and electro-acupuncture in target points and organs (Appendix S8), future studies in either field may benefit from cross-referencing literature in the other field.

A major limitation of this study was the confounding effects of anesthesia during PTNS, which can lead to physiological changes, as well as animal death. Anesthesia allows for measurement of vaginal blood perfusion and accurate sustained placement of the stimulation wire. However, other studies that saw an increase in serum estradiol levels did not anesthetize their animals [13, 15, 16]. Ketamine-xylazine anesthesia has been demonstrated to reduce nerve conduction velocities [37] and cystometric responses [38] in mice. In addition to local interference at the nerve level, ketamine-xylazine reduces glucose metabolism in the brain [39], a site of extragonadal aromatization. Ketamine-xylazine anesthesia has also resulted in delayed recovery and possible hypoxia [40]. Animal death secondary to anesthesia decreased the sample size and strength of this study and may have contributed to variability in group measures. In this study, many comparisons between OVX stim and control groups were close to significance (p = 0.05 – 0.1), and more definitive conclusions may have been drawn without animal loss decreasing sample size. Our mortality rate of 2% is similar to previously documented mortality rates (2.01%) in rats [41]. Other limitations include the fact that the OVX rat model we used differs from natural human menopause (Appendix S9) and the age for performing the OVX surgery ranges (Appendix S10) depending on the study objectives. Future studies may consider stimulation of awake animals or use of different OVX models to combat these confounders.

The expansion of PTNS beyond traditional target organs such as the bladder improves our understanding of its mechanism of action. The positive effect of PTNS on vaginal blood perfusion is consistent with the current hypothesis of how PTNS works (stimulation of afferent pathways to the lumbosacral spinal cord leading to modulation of efferent outflow to corresponding pelvic organs), since the bladder and vagina have shared innervation (pudendal and pelvic nerves) [42] and are physically far from the site of stimulation (ankle). The lack of response in vaginal blood perfusion after week 5 of the experiment suggests PTNS is limited to organs that have adequate innervation and vascular structure. However, unilateral findings (positive effect only on the side of stimulation) in osteoporotic parameters (bone mineral density and yield load) suggest that part of the effect may be local. Indeed, neuromuscular electrical stimulation may stimulate osteogenesis by applying mechanical stress to the bone via muscle contractions [43]. Additional studies to elucidate the full potential of PTNS on menopause-associated systems under different experimental conditions are warranted.

## CONCLUSION

In this preliminary study, PTNS did not increase serum estradiol levels, but transiently increased VBP during stimulation for up to 5 weeks after OVX and increased aBMD and yield load compared to unstimulated OVX controls independent of estradiol levels. Therefore, PTNS may ameliorate some symptoms associated with menopause. However, this study had a variety of limitations including small animal numbers due to animal loss and the confounding effects of anesthesia. Previous literature suggests that electro-acupuncture at the “SP6” acupoint increased serum estradiol levels in both rats and humans [13, 14]. Future studies may include 1) modifying stimulation frequency and duration within and between sessions, 2) varying points of stimulation (unilateral, bilateral, or in conjunction with other sites), 3) using a different model of menopause (naturally aged or VCD-induced), and 4) conducting awake rat studies to eliminate anesthetic-related confounders (implantable nerve cuff, habituation of rat to restraint and stimulation, introduction of wire under temporary anesthesia followed by awake stimulation).

## ACKNOWLEDGEMENTS

We thank Teera Losch from the U-M Core Assay Facility lab for her assistance in estradiol kit selection and sample processing, Yesen Zhou and Christopher Fry in Jean Nemzek’s lab for their assistance with processing and storing samples, Ingrid Bergin from U-M In-Vivo Animal Core for her assistance with histology and expertise on the uterotrophic assay, Dana Jackson for his assistance with preliminary biomechanical testing, Jill Becker for her advice during experimental design, and the Unit for Laboratory Animal Medicine for their care of the animals. We would like to acknowledge support from the National Institute of Arthritis and Musculoskeletal and Skin Diseases for the Michigan Integrative Musculoskeletal Health Core Center (P30 AR069620).

