## Supplementary Information for "Tibial nerve stimulation increases vaginal blood perfusion and bone mineral density and yield load in ovariectomized rat menopause model"

**Appendix contents:** Appendix items shown according to the section of the paper they are referenced in.


1. Introduction
2. Methods
    a. Appendix S1: OVX and sham procedures
    b. Appendix S2: Anesthetic regime modification
    c. Appendix S3: Percutaneous tibial nerve stimulation
    d. Appendix S4: Post mortem femur analysis, micro-CT reconstruction
3. Results
    a. Appendix S5: Animal loss during study
    b. Appendix S6: Blood glucose, serum estradiol, vaginal cytology
4. Discussion
    a. Appendix S7: Proposed mechanisms of electro-acupuncture on extragonodal aromatization of estradiol in an ovariectomized rat
    b. Appendix S8: Relationship between electro-acupuncture and neuromodulation fields
    c. Appendix S9: Rat models of menopause
    d. Appendix S10: Age for performing OVX surgery in rats
5. Conclusion


**Appendix S1: OVX and sham procedures**:

**OVX procedure**: The OVX was performed with a dorsal skin incision, consistent with previously described methods [1]. On each side, the abdominal wall was punctured below the level of the ribcage with a stab incision. The ovary was located through the incision and removed from the uterine horn through tearing of the tissue between two clamped hemostats. The hemostats were kept on the pedicle for 30 seconds, and hemostasis was confirmed before the uterus was dropped back into the abdominal cavity. The puncture in the abdominal wall was closed with simple interrupted absorbable sutures. After both sides of the abdominal wall were closed, the skin was closed with wound clips.

**Sham procedure**: The same surgical procedures were performed as the OVX surgery up to the point of abdominal incision, but the ovaries were not manipulated or removed, and the abdominal wall was sutured closed immediately after the incisions were made. The surgical closure was the same as for the OVX groups.

**Appendix S2: Anesthetic regime modification:** The reversal agent atipamezole was added after the first week of the study to improve recovery time. Subcutaneous (compared to intraperitoneal) ketamine-xylazine was trialed in 3 rats (1 from each test group) during one session. However, rats did not reach appropriate plane of anesthesia, and intraperitoneal injection was resumed for the remainder of the study. During weeks 1 and 2, rats were re-dosed with ketamine-xylazine on the same day if they did not reach an appropriate plane of anesthesia. After week 3, the rats were recovered and were re-dosed the following day to prevent anesthetic-related death.

**Appendix S3: Percutaneous tibial nerve stimulation**: These settings were selected based on previous human [2] and rat [3] tibial nerve stimulation studies in our laboratory. The right ankle was preferentially simulated, but when unsuccessful, the left side was attempted. During week 1 and week 2, four rats and one rat respectively had their left side stimulated after right side wire placement was unsuccessful. The remainder of the stimulations were performed successfully on the right side.

**Appendix S4: Post mortem femur analysis, micro-CT reconstruction**: Femurs were scanned at 9 µm isotropic voxel size with a 0.3° rotation angle, 2 frames averaged, a 1 mm aluminum filter, with a source voltage of 70 kVp, and source current of 357 µA. Micro-CT images were then reconstructed and calibrated to manufacturer supplied phantoms of hydroxyapatite. In the femora, regions of interest (ROI) were created for both cortical and trabecular bone regions. The cortical ROI spanned 15% of total bone length centered between the proximal end of the distal femoral growth plate and the distal end of the third trochanter. To segment the cortical bone, Ostu threshold values were generated for each sample, and a fixed value (572 mg/cm$^3$) representing the mean of all samples was selected for all groups. CTAnalysis (Skyscan 1176, Bruker, Billerica, MA) was used to measure total area, bone area, marrow area, cortical thickness, endosteal perimeter, periosteal perimeter, anterior-posterior bending moment of inertia, and tissue mineral density (TMD). The trabecular region of interest (ROI) spanned 10% of the bone length starting at the proximal end of the distal femoral growth plate and extending proximally into the metaphysis. Trabecular bone was segmented using a two-phase adaptive threshold available through CTAnalysis. An initial pre-threshold of 376 mg/cm$^3$ was used, which was calculated from the average Otsu threshold for the ROIs. The samples were then more accurately segmented via the adaptive mean of minimum and maximum threshold values within a five-pixel radius of the target pixels available through CTAnalysis.

**Appendix S5: Animal loss during study**: One animal was euthanized due to a post-surgical complication before the start of the study (OVX stim group). Of the remaining twenty-nine animals, six animals were lost during anesthetic events, usually experiencing prolonged recovery after anesthesia, and expiring during the recovery process. While most animals recovered within 30-60 minutes of the atipamazole reversal, several animals remained lethargic for upwards of 2 hours even with reversal. Two animals were lost (both OVX control group) during week 1, two animals (both OVX stim group) during week 3, one animal (OVX control) during week 4, and one animal (intact control) during the first half of week 6. Finally, one animal (OVX control) died during second sedated session of week 6 but was necropsied immediately upon death and counted within the necropsy results.

**Appendix S6: Blood glucose, serum estradiol, vaginal cytology**: Blood glucose measures ranged from 81-185 mg/dL across all animals and sessions. No significant differences were found between groups or over time. Starting from week 3 of the experiments (5 weeks after OVX surgery), the OVX stim and OVX control groups had significantly decreased serum estradiol measures compared to the intact control group, except for week 5 (OVX control vs intact control not different), week 6 (OVX control vs intact control not different), and at necropsy (OVX stim and intact control, not different). The serum estradiol for OVX control and OVX stim groups were not significantly different at any time point (Table S4). Vaginal cytology for all quantified weeks (weeks 3, 4, 6) differed significantly between the intact control group and the OVX control and OVX stim groups but was not significantly different between the two OVX groups. The intact control group had a greater percent of cornified and epithelial cells while the OVX control and OVX stim had a greater fraction of leukocytes (leukocytes, cornified cells, epithelial cells respectively: intact control = 0.2 ± 0.3, 0.8 ± 0.2, 0.8 ± 0.2; OVX control = 0.4 ± 0.3, 0.1 ± 0.1, 0.1 ± 0.1; OVX stim = 0.3 ± 0.02, 0.05 ± 0.05, 0.06 ± 0.08).

**Appendix S7: Proposed mechanisms of electro-acupuncture on extragonodal aromatization of estradiol in an ovariectomized rat.** The exact mechanisms of the effects of dorsal and abdominal midline acupoints are unknown, but may include stimulation of ovarian and ovarian plexus nerves involved in the modulation of ovarian estradiol secretion (in intact animals) [4], stimulation of the estradiol precursors such as androstendione in adrenal cortex by the splanchnic nerve [5], and stimulation at extragonodal sites of aromatization (conversion to estrogens) such as the mesenchymal cells of adipose tissue, osteoblasts and chondrocytes of the bone, vascular endothelium and aortic smooth muscle cells, and numerous sites in the brain [6].

**Appendix S8: Relationship between electro-acupuncture and neuromodulation fields**: A major challenge in interpreting this study is the lack of communication between the electro-acupuncture and neuromodulation fields in the literature. Tibial nerve stimulation in the neuromodulation field primarily focuses on pelvic organs such as the bladder or bowel, with limited literature on the reproductive tract. In contrast, electro-acupuncture studies primarily focus on stimulating the "SP6" acupoint alone or in conjunction with other acupoints to treat hormone-associated disorders such as menopause [7], primary dysmenorrhea (menstrual cramps) [8, 9] and labor [10–12]. However, electro-acupuncture has also been indicated in the treatment of urinary (overactive bladder [13], stress urinary incontinence [14]) or GI disease (fecal incontinence [15]). Despite the significant overlap between tibial nerve stimulation and electro-acupuncture at the "SP6" acupoint in method and target organ effect, only a few sources [16] cross-referenced or acknowledged similar studies in the corresponding field. This is likely because the connection between the two fields is not intuitive, and there may be language barriers. We strongly encourage future studies to search both electro-acupuncture and neuromodulation literature to better understand and interpret from the existing body of literature.

**Appendix S9: Rat models of menopause**: Compared to aged rats that naturally reach estropause at 9-12 months (rat equivalent of menopause, both a persistent stage of reproductive senescence), the OVX rat model has a different hormone profile and hypothalamic–pituitary–gonadal axis compared to animals with an intact reproductive tract [22]. However, aged rats in estropause differ from the human menopause condition in that they remain in persistent estrus (characterized by moderate to high estradiol levels) compared to human menopause, which is characterized by low or undetectable levels of estradiol. While both aged rats in estropause [22] [23], and the surgically induced OVX model [19] have been previously used to demonstrate the effect of PTNS stimulation on increasing serum estradiol levels, an alternative model of transition menopause induced by the drug 4-vinylcyclohexene diepoxide (VCD) could more closely replicate the human condition. The VCD-induced model depletes the resting follicle pool but allows for the retention of follicle-deplete ovarian tissue, resulting in an ovarian and hormone profile more similar to women undergoing natural menopause [22].

**Appendix S10: Age for performing OVX surgery in rats**: The age for performing OVX surgery varies from around 12 weeks of age at sexual maturity (as in this study) [24], 6 months for models of osteoporosis [1], to 8 months for models of postmenopausal hypertension [25]. In this study 250-300 g rats (11 – 15 weeks) were chosen based on previous studies of tibial nerve stimulation to drive genital arousal [3] [26].

**Table S1:** p values of Tukey's HSD for normalized weight. Significance: * $p < 0.05$, ** $p < 0.01$. NS = no significance.

| Normalized weight | OVX control vs intact control (▲) | OVX stim vs intact control (●) | OVX control vs OVX stim |
|---|---|---|---|
| 1 week post sx | 0.0199* | NS | NS |
| W1 | 0.0041** | 0.0002** | NS |
| W2 | 0.0695 | 0.0398* | NS |
| W3 | 0.0004 ** | < 0.0001** | NS |
| W4 | NS | 0.0326* | NS |
| W5 | 0.0011** | 0.0544 | NS |
| W6 | 0.0017** | 0.0299* | NS |

| | | | |
|---|---|---|---|
| Necropsy | 0.0038** | 0.0733 | NS |

**Table S2**: p values of Wilcoxon signed-rank test (probability < t =1), comparing normalized vaginal blood perfusion at each timepoint to the expected value of 1. Significance: * p < 0.05, ** p < 0.01. NS = no significance.

| | Timepoint | Intact control | OVX control | OVX stim |
|---|---|---|---|---|
| Week 1 | 0 - 10 min | NS | NS | 0.0039** |
| | 10 – 20 min | NS | NS | 0.0020** |
| | 20 – 30 min | 0.0195* | NS | 0.0020** |
| Week 3 | 0 - 10 min | NS | NS | NS |
| | 10 – 20 min | NS | NS | 0.0391* |
| | 20 – 30 min | NS | NS | 0.0117* |
| Week 6 | 0 - 10 min | 0.0488* | NS | NS |
| | 10 – 20 min | NS | NS | NS |
| | 20 – 30 min | NS | NS | NS |

**Table S3**: p values of Steel-Dwaas test of normalized 10-min bins of vaginal blood perfusion (VBP), binned in 10 minute intervals, comparing between animal groups. Significance: * p < 0.05, ** p < 0.01. NS = no significance.

| VBP week | Timepoint | OVX control vs intact control (▲) | OVX stim vs intact control (●) | OVX control vs OVX stim (*) |
|---|---|---|---|---|
| W1 | 0-10 min | NS | 0.0220* | NS |
| | 10-20 min | NS | 0.0282* | NS |
| | 20-30 min | NS | NS | NS |
| W3 | 0-10 min | NS | NS | NS |
| | 10-20 min | NS | NS | NS |
| | 20-30 min | NS | NS | NS |
| W6 | 0-10 min | NS | NS | NS |
| | 10-20 min | NS | NS | NS |
| | 20-30 min | NS | NS | NS |

**Table S4**: p values of Tukey's HSD for serum estradiol. Significance: * p < 0.05, ** p < 0.01. NS = no significance.

| Serum estradiol | OVX cont vs intact control (▲) | OVX stim vs intact control (●) | OVX control vs OVX stim |
|---|---|---|---|
| Sx | NS | NS | NS |
| W1 | NS | NS | NS |
| W2 | NS | NS | NS |
| W3 | 0.0293* | 0.0216* | NS |
| W4 | 0.0071** | 0.0023** | NS |
| W5 | NS | 0.0037** | NS |
| W6 | NS | 0.0069** | NS |
| Necropsy | 0.0344* | NS | NS |

**Table S5**: Values from DXA scanning, reported as mean ± standard deviation.

| | Intact control | OVX control | OVX stim |
|---|---|---|---|
| R aBMD (g/cm$^2$) | 0.20 ± 0.008 | 0.18 ± 0.009 | 0.19 ± 0.01 |
| R BMC (g) | 0.45 ± 0.03 | 0.40 ± 0.03 | 0.43 ± 0.04 |
| R BA (cm$^2$) | 2.24 ± 0.09 | 2.24 ± 0.14 | 2.28 ± 0.12 |
| L aBMD (g/cm$^2$) | 0.20 ± 0.008 | 0.18 ± 0.009 | 0.19 ± 0.01 |
| L BMC (g) | 0.45 ± 0.03 | 0.40 ± 0.009 | 0.44 ± 0.04 |
| L BA (cm$^2$) | 2.21 ± 0.03 | 2.18 ± 0.06 | 2.32 ± 0.10 |

**Table S6**: Values from micro-CT analysis of right femurs, reported as mean ± standard deviation. (* $p < 0.05$ vs. Intact control).

| | Intact control | OVX control | OVX stim |
|---|---|---|---|
| Trabecular bone mineral density (BMD) (g/cm$^3$) | 0.42 ± 0.06 | 0.25 ± 0.05 * | 0.30 ± 0.07 * |
| Trabecular bone volume fraction (BV/TV, %) | 33.80 ± 5.00 | 20.46 ± 3.70 * | 23.46 ± 3.25 * |
| Trabecular thickness (Tb.Th, mm) | 0.08 ± 0.003 | 0.09 ± 0.002 | 0.09 ± 0.003 |
| Trabecular number (Tb.N, 1/mm) | 4.08 ± 0.54 | 2.34 ± 0.40 * | 2.67 ± 0.36 * |
| Cortical tissue mineral density (TMD) (g/cm$^3$) | 0.19 ± 0.08 | 1.22 ± 0.08 | 1.22 ± 0.08 |
| Cortical total area (mm$^2$) | 9.84 ± 0.90 | 9.41 ± 0.64 | 10.09 ± 0.94 |
| Cortical bone area (mm$^2$) | 5.96 ± 0.40 | 5.50 ± 0.24 | 5.81 ± 0.53 |
| Cortial cross sectional thickness (mm) | 0.46 ± 0.09 | 0.53 ± 0.03 | 0.51 ± 0.04 |
| Polar moment of inertia (mm$^4$) | 4.49 ± 0.69 | 3.86 ± 0.51 | 4.54 ± 1.02 |
| Bone marrow area (mm$^2$) | 3.90 ± 0.58 | 3.91 ± 0.63 | 4.27 ± 0.59 |

**Table S7**: Values from biomechanical testing, reported as mean ± standard deviation. (* $p < 0.05$ vs. OVX control)

| | Intact control | OVX control | OVX stim |
|---|---|---|---|
| Stiffness (N/mm) | 545.34 ± 51.19 | 511.67 ± 89.00 | 580.90 ± 88.50 |
| Yield load (N) | 145.72 ± 11.62 | 132.46 ± 10.76 | 152.07 ± 16.36 * |
| Ultimate load (N) | 186.88 ± 18.27 | 181.13 ± 8.18 | 199.88 ± 28.77 |
| Fail load (mm) | 175.11 ± 22.66 | 159.33 ± 29.45 | 192.75 ± 25.54 |

| | | | |
|---|---|---|---|
| Yield displacement (mm) | 0.28 ± 0.03 | 0.28 ± 0.07 | 0.23 ± 0.08 |
| Ultimate displacement (mm) | 0.58 ± 0.08 | 0.64 ± 0.12 | 0.61 ± 0.07 |
| Fail displacement (mm) | 0.69 ± 0.18 | 0.92 ± 0.20 | 0.80 ± 0.19 |
| Total work (Nmm) | 95.27 ± 34.55 | 125.74 ± 33.30 | 115.55 ± 39.88 |
| Post yield displacement (mm) | 0.42 ± 0.19 | 0.64 ± 0.18 | 0.52 ± 0.19 |

**Table S8:** p values of Tukey's HSD for femur analysis. Significance: * p < 0.05, ** p < 0.01. NS = no significance.

| Mode of analysis | | OVX control vs intact control | OVX stim vs intact control | OVX control vs OVX stim |
|---|---|---|---|---|
| DXA | Right femur areal bone mineral density (aBMD) | 0.0002** | 0.0908 | 0.0463* |
| | Left femur aBMD | 0.0014* | 0.0669 | NS |
| Micro-CT (Trabecular) | BMD (g/cm$^3$) | <0.0001** | 0.0022** | NS |
| | Bone volume fraction | <0.0001** | 0.0002** | NS |
| | Thickness | 0.0075** | 0.0024** | NS |
| | Number | <0.0001** | <0.0001** | NS |
| Micro-CT (Cortical) | Tissue mineral density (TMD) | NS | NS | NS |
| | Total area | NS | NS | NS |
| | Bone area | NS | NS | NS |
| | Cross-sectional thickness | NS | NS | NS |
| Micro-CT (Other) | Moment of inertia | NS | NS | NS |
| | Marrow area | NS | NS | NS |
| Biomechanical | Stiffness | NS | NS | NS |
| | Yield Load | NS | NS | 0.0272* |
| | Ultimate load | NS | NS | NS |
| | Fail load | NS | NS | NS |
| | Yield displacement | NS | NS | NS |
| | Ultimate displacement | NS | NS | NS |
| | Fail displacement | NS | NS | NS |
| | Total work | NS | NS | NS |
| | Post yield displacement | NS | NS | NS |

**Supplementary Information References**:

13. Paik, S.-H., Han, S.-R., Kwon, O.-J., Ahn, Y.-M., Lee, B.-C., & Ahn, S.-Y. (2013). Acupuncture for the

treatment of urinary incontinence: A review of randomized controlled trials. *Experimental and therapeutic medicine*, *6*(3), 773–780. https://doi.org/10.3892/etm.2013.1210

26. Zimmerman, L. L., Mentzelopoulos, G., Parrish, H. J., Luma, B. D., Becker, J. B. Bruns, T. M. (2019).

Modulating sexual behavior in female rats with tibial nerve electrical stimulation. Society for Neuroscience Annual Meeting [Abstract]. Retrieved from https://www.abstractsonline.com/pp8/#!/7883/presentation/70326
